# Horizontal gene transfer and CRISPR targeting drive phage-bacterial host interactions and co-evolution in pink berry marine microbial aggregates

**DOI:** 10.1101/2023.02.06.527410

**Authors:** James C. Kosmopoulos, Danielle E. Campbell, Rachel J. Whitaker, Elizabeth G. Wilbanks

## Abstract

Bacteriophages (phages), viruses that infect bacteria, are the most abundant components of microbial communities and play roles in community dynamics and host evolution. The study of phage-host interactions, however, is made difficult by a paucity of model systems from natural environments and known and cultivable phage-host pairs. Here, we investigate phage-host interactions in the “pink berry” consortia, naturally-occurring, low-diversity, macroscopic aggregates of bacteria found in the Sippewissett Salt Marsh (Falmouth, MA, USA). We leverage metagenomic sequence data and a comparative genomics approach to identify eight compete phage genomes, infer their bacterial hosts from host-encoded clustered regularly interspaced short palindromic repeats (CRISPR), and observe the potential evolutionary consequences of these interactions. Seven of the eight phages identified infect the known pink berry symbionts *Desulfofustis* sp. PB-SRB1, *Thiohalocapsa* sp. PB-PSB1, and *Rhodobacteraceae* sp. A2, and belong to entirely novel viral taxa, except for one genome which represents the second member of the *Knuthellervirus* genus. We further observed increased nucleotide variation over a region of a conserved phage capsid gene that is commonly targeted by host CRISPR systems, suggesting that CRISPRs may drive phage evolution in pink berries. Finally, we identified a predicted phage lysin gene that was horizontally transferred to its bacterial host, potentially via a transposon intermediary, emphasizing the role of phages in bacterial evolution in pink berries. Taken together, our results demonstrate that pink berry consortia contain diverse and variable phages, and provide evidence for phage-host co-evolution via multiple mechanisms in a natural microbial system.

**IMPORTANCE:** Phages (viruses that infect bacteria) are important components of all microbial systems, where they drive the turnover of organic matter by lysing host cells, facilitate horizontal gene transfer (HGT), and co-evolve with their bacterial hosts. Bacteria resist phage infection, which is often costly or lethal, through a diversity of mechanisms. One of these mechanisms are CRISPR systems, which encode arrays of phage-derived sequences from past infections to block subsequent infection with related phages. Here, we investigate bacteria and phage populations from a simple marine microbial community known as “pink berries” found in salt marshes of Falmouth, Massachusetts, as a model of phage-host co-evolution. We identify eight novel phages, and characterize a case of putative CRISPR-driven phage evolution and an instance of HGT between phage and host, together suggesting that phages have large evolutionary impacts in a naturally-occuring microbial community.

## INTRODUCTION

Phages, viruses that infect bacteria, occur in all microbial ecosystems, often outnumbering bacteria by 10 to 1, and play pivotal roles in altering community structure (Andersson & Banfield, 2008; Bergh *et al*., 1989; Breitbart *et al*., 2018), mediating horizontal gene transfer (HGT) (Breitbart *et al*., 2018; Hall *et al*., 2017; Schneider, 2021), and driving bacterial evolution (Campbell *et al*., 2020; Koskella & Brockhurst, 2014; Martiny *et al*., 2014). Though some phage-host interactions can be beneficial, phage infection canonically ends with the lysis and death of the host to release progeny phage particles for transmission to new host cells. Thus, there is strong selection for bacteria to evolve mechanisms to resist infection. Likewise, phages must evolve to overcome those resistances to survive. This phage-bacterial host coevolution is often described as an “arms race” (Hampton *et al*., 2020).

Bacteria have evolved a wide range of phage defense systems, such as clustered regularly interspaced short palindromic repeat (CRISPR) loci, which act as microbial adaptive immune systems. During a new phage infection, CRISPR systems incorporate short segments of phage-derived sequence, known as “protospacers,” into CRISPR arrays as “spacers” (Barrangou *et al*., 2007; Jansen *et al*., 2002; Jinek *et al*., 2012). CRISPR systems further encode mechanisms to degrade invading phage DNA that matches an existing spacer, allowing the host to resist infection. CRISPR arrays thus serve as a genetic record of phages a host has encountered and can be leveraged to identify phage hosts from sequence data (Childs *et al*., 2014; England *et al*., 2018).

In-depth analysis of CRISPR targeting offers insights into phage-host interactions. CRISPR systems often target conserved phage sequences, thus conferring protection from groups of related phages (Barrangou *et al*., 2007; Deveau *et al*., 2008; Mojica *et al*., 2009). Phages can accumulate mutations within protospacers which allow the phage to escape CRISPR defenses (Andersson & Banfield, 2008; Deveau *et al*., 2008; Sun *et al*., 2013). Together, these patterns in CRISPR spacer and protospacer nucleotide variation can shed light on phage-host co-evolution.

Here, we investigate phage-bacterial interactions in microbial consortia known as “pink berries.” Pink berries are macroscopic microbial aggregates found in the Sippewissett Salt Marsh of Falmouth, MA (Seitz *et al*., 1993; Wilbanks *et al*., 2014). These aggregates are primarily composed of phototrophic, sulfide-oxidizing and sulfate-reducing bacteria that together form a syntrophic sulfur cycle (Wilbanks *et al*., 2014). Though phage sequences have been found in pink berries previously (Wilbanks *et al*., 2022), the interactions of phage and their host remain uncharacterized. We show that pink berries host diverse, novel phages that drive bacterial evolution through HGT and putative CRISPR-driven arms race dynamics.

## RESULTS

### Pink berries contain novel phages that are variable between individual aggregates

Pink berries were strategically sampled and pooled during sequencing library preparation to observe both the breadth of diversity across individual pink berries, and to deeply sample the total diversity of pink berries. Thus, we sequenced two metagenomes from single pink berries, one with low sequencing depth (LS06-2018-s01) and one with high sequencing depth (LS06-2018-s02), and one metagenome from three pink berries homogenized together and sequenced at very high depth (LS06-2018-s03) (Table 1). Co-assembly of the three pink berry metagenomes yielded 184 contigs totaling 4.35 Mb in length (Table 1). Two phage sequence prediction tools, VIBRANT (Kieft *et al*., 2020) and ViralVerify (Antipov *et al*., 2020), identified nine full-length, circular phage genomes which were the targets of our downstream analyses (Suppl. Fig. 1). The phages were named according to their hosts predicted by CRISPR spacer-protospacer matches (described in *Pink berry phages are targeted by bacterial CRISPR systems)*.

**Table 1.**
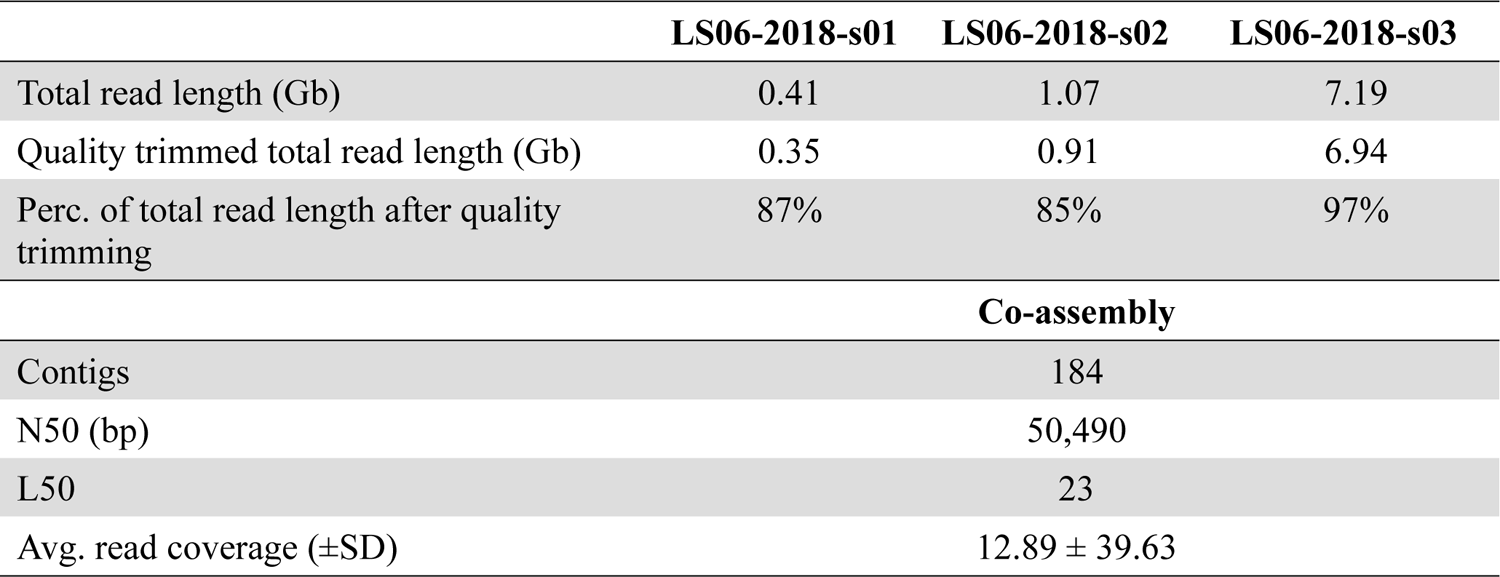
Summary of pink berry metagenome sequencing and co-assembly.

**Figure 1.**
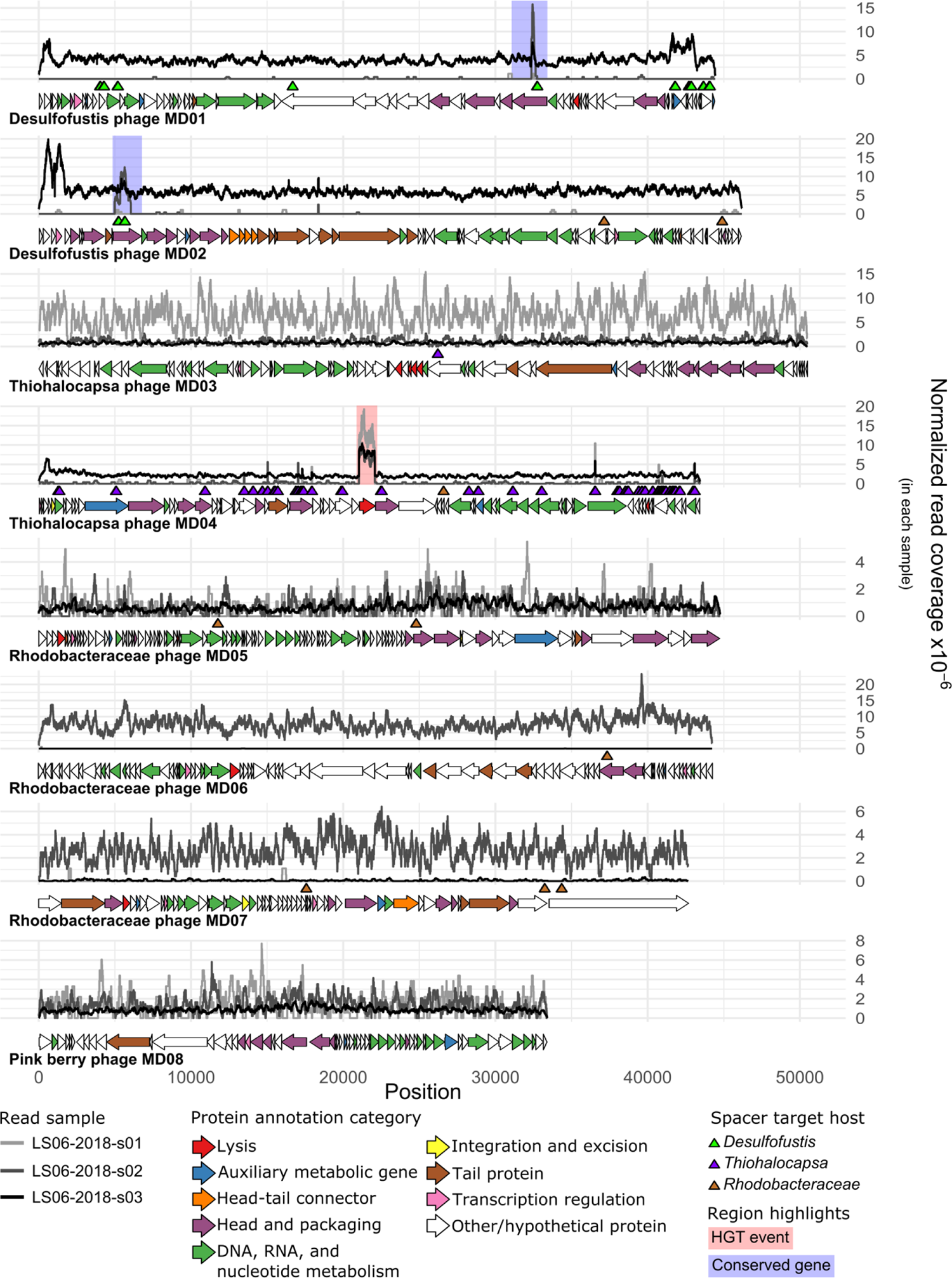
Complete phage genomes vary in abundance across samples and are targeted by bacterial CRISPR spacers. Normalized read coverage by position for each sample are given. Coverage values were normalized to the total number of trimmed and filtered reads for each sample. Horizontal arrows indicate ORFs predicted by PHANOTATE (McNair *et al*., 2019), and their colors correspond to predicted functional categories. Triangles indicate genome positions of protospacers, colored by host taxonomy of the corresponding spacer. Regions highlighted with a blue background indicate a conserved gene between phage genomes inferred by Clinker (Gilchrist & Chooi, 2021). Region highlighted with a red background was found to be an HGT event between the phage and host.

Functional annotations were obtained for 236 of 632 putative protein-coding genes predicted across the nine complete viral genomes (Suppl. Data 1). One virus was found to have large amounts of homology to eukaryote-infecting Circular Rep-encoding Single Stranded (CRESS) DNA viruses and may infect a nematode host, which are frequently observed grazing on bacteria in pink berries. This virus was excluded from further analysis to focus on the primary pink berry bacterial components and their phages (“Pink berry virus MD00”; Suppl. Data 1). Functional annotations for the remaining 8 phage genomes predicted nucleotide metabolism proteins, head and packaging proteins, integration and excision proteins, transcriptional regulators, and various lytic proteins (Fig. 1, Suppl. Data 1). Importantly, all phages of interest here are predicted to exhibit strictly lytic lifecycles, as they lack the genes for a temperate lifecycle, such as an integrase. There were also predictions for several auxiliary metabolic genes (Breitbart *et al*., 2007), which we inferred based on functional predictions outside the core functions required for the phage lifecycle. One such AMG was a *darB*-like antirestriction gene encoded on the Thiohalocapsa phage MD04 genome (Suppl. Data 1). Interestingly, *darB* has been shown to methylate phage DNA to resist host restriction modification (RM) systems (Iida *et al*., 1987; Iyer *et al*., 2017). Prior work found pink berry bacteria employ numerous, diverse RM systems, and revealed that putative pink berry phages have been shown to contain similar methylation profiles as their hosts (Wilbanks *et al*., 2022). The presence of AMGs such as *darB* suggest that pink berry phages have adapted to increase their fitness in *Thiohalocapsa* hosts, consistent with an armsrace-like process of coevolution.

Phage taxonomy is based on genome similarity (Turner *et al*., 2021), and forms the basis of inferring the diversity of phages within a community. We hypothesized that pink berry phages would be at least as diverse as their pool of potential bacterial hosts, which span multiple phyla. vConTACT (Bin Jang, *et al*., 2019), a genome-wide protein similarity-based approach, was used to infer phage taxonomy for the eight phage genomes of interest. Although nominal protein similarity was detected between all phages of interest and the Viral RefSeq protein database during gene annotation (Suppl. Data 1), only Desulfofustis phage MD02, Thiohalocapsa phage MD04, and Rhodobacteraceae phage MD07 are connected to a known phage in the vConTACT network (Fig. 2, Suppl. Data 2). Further, only one phage genome, Desulfofustis phage MD02, had sufficient protein similarity to cluster with a cultured phage reference genome, Pseudomonas phage PMBT14, which is currently the only species of the genus *Knuthellervirus* (Suppl. Data 2). None of the phage genomes of interest clustered with each other. The remaining five genomes of interest lacked sufficient protein similarity for a connection in the vConTACT network, indicating that these phages represent novel and undescribed diversity.

**Figure 2.**
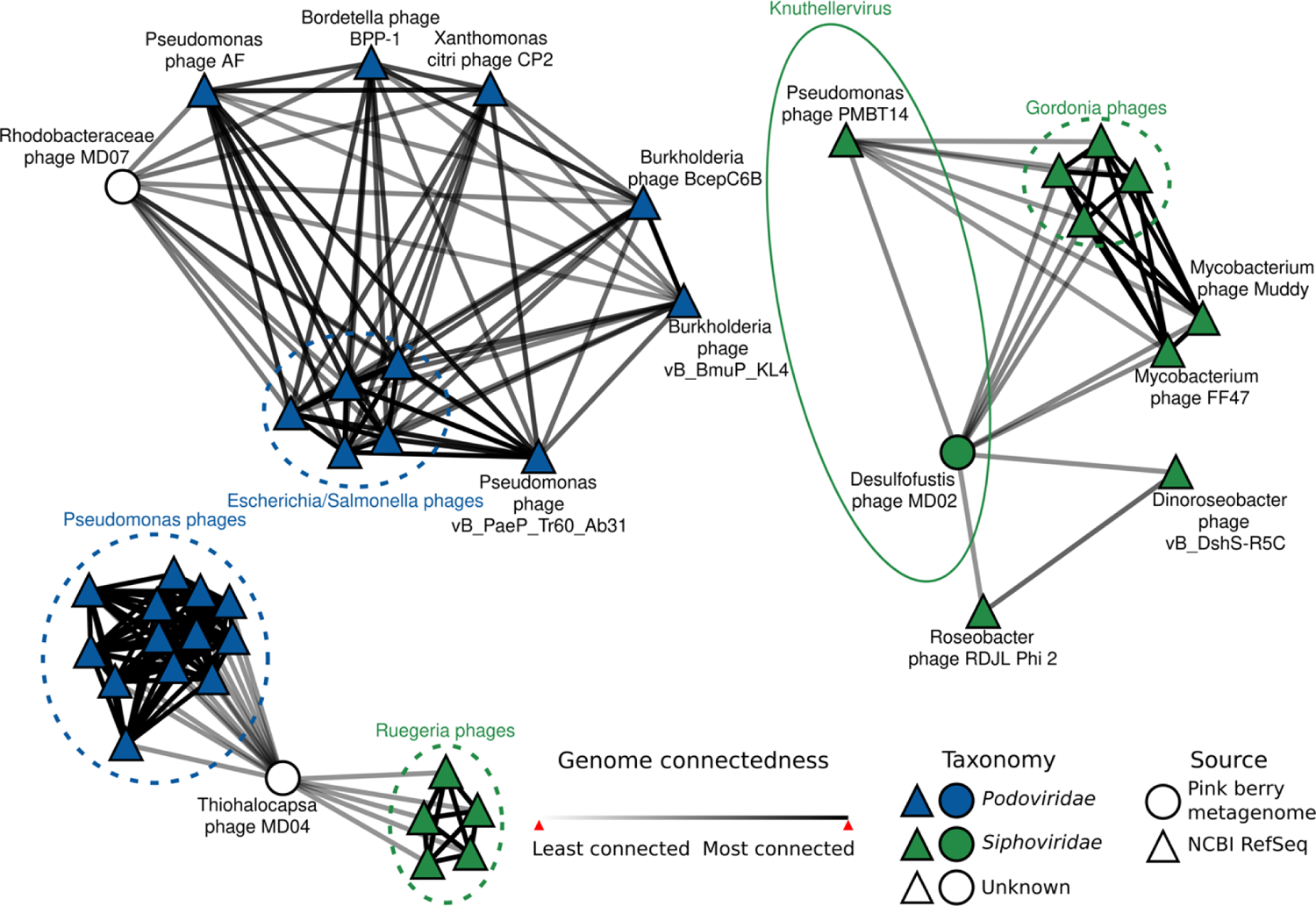
Pink berry consortia host largely novel phages. Each node represents a phage genome and edges are genome relatedness inferred by vConTACT (Bin Jang *et al*., 2019). Reference genomes are from the RefSeq Prokaryotic virus database v211 (Brister *et al*., 2015). The group circled with a solid line shows the phage genus of interest, *Knuthellervirus*. Groups circled in dashed lines show related phages infecting the same host. The network was visualized in Cytoscape v3.9.0 (Su *et al*., 2014). Only phages which are first-neighbors to pink berry phage genomes are shown.

Relative bacterial abundances are similar across pink berries (Wilbanks *et al*., 2014), which is likely due to the constraints of the syntrophic metabolic interactions between constituent bacteria. To assess the distribution of pink berry-associated bacteria and phages, genome-wide read coverages were analyzed for individual pink berry metagenomes (Fig. 3). In agreement with previous observations, we found that the relative abundance of pink berry bacteria is relatively homogenous across samples (Fig. 3A). In contrast, phage presence and abundance are highly variable between different pink berry communities (Fig. 3B). Only two phages, Rhodobacteraceae phage MD05 and Pink berry phage MD08, are similarly abundant in each pink berry metagenome, while read mapping to the remaining six phages suggests they are distributed unevenly between individual aggregates (Fig. 3). Additionally, to observe whether the phage genomes of interest are present in pink berry metagenomes from previous years, we mapped reads from a metagenome of 10 pink berries sampled in 2011 (Wilbanks *et al*., 2014). This revealed near-complete coverage of Desulfofustis phage MD01 and Rhodobacteraceae phage MD06 (Suppl. Fig. 2), indicating that these phages have persisted over seven years. In contrast, read mapping to Desulfofustis phage MD02 and Thiohalocapsa phage MD04 genome largely occurs at highly conserved regions, and is likely the consequence of non-specific cross-mapping (Suppl. Fig. 2). Likewise, Thiohalocapsa phage MD03, Rhodobacteraceae phages MD05 and MD07, and Pink berry phage MD00 had <1% or no genome coverage. Since neither this study nor the study from 2011 (Wilbanks *et al*., 2014) are exhaustive surveys of pink berry diversity, it is difficult to determine the mechanisms behind the emergence of these six phages. Taken together, these results suggest that although pink berries have relatively simple and conserved bacterial community structures, their phages are highly variable over both space and time.

**Figure 3.**
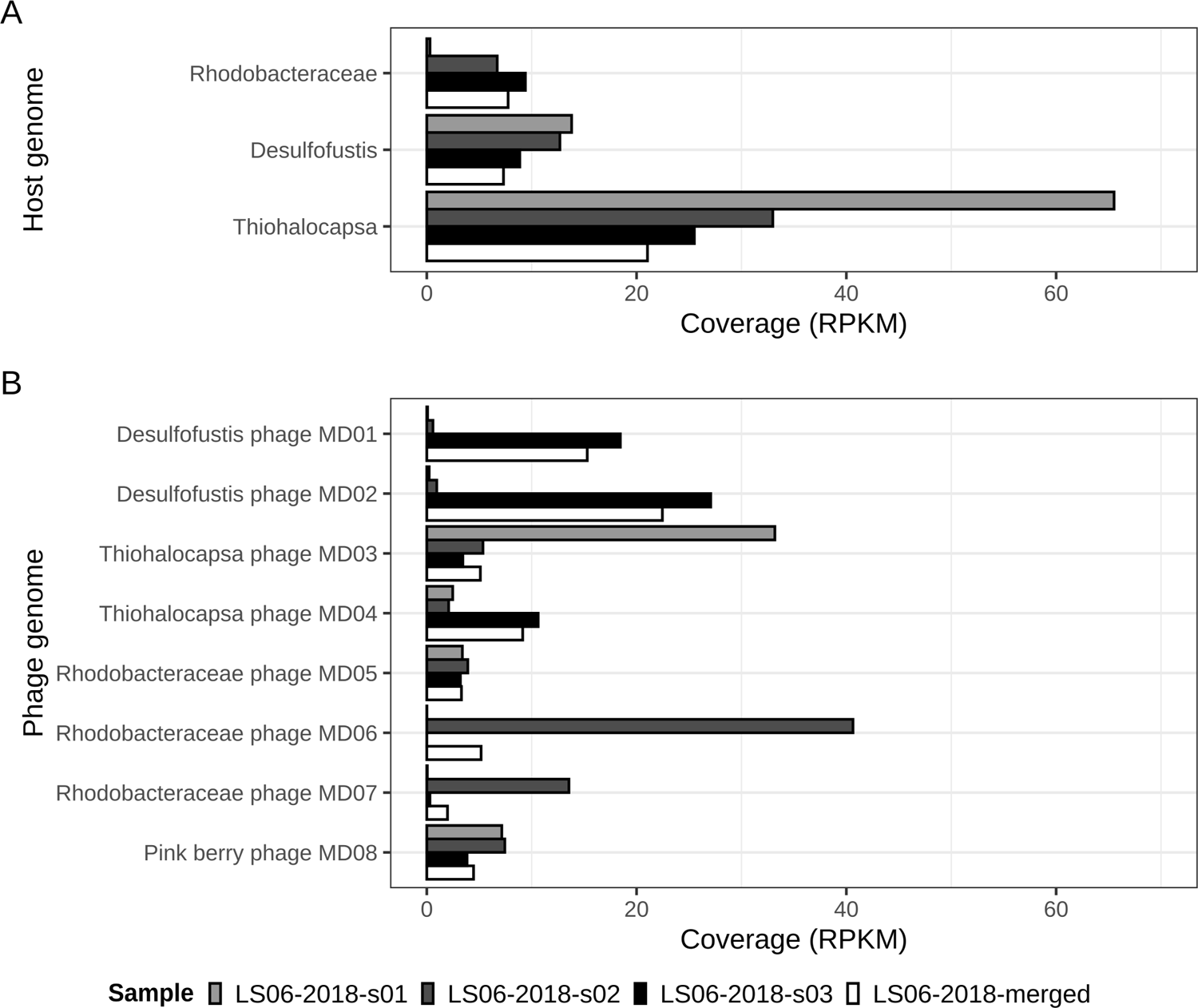
Phage presence and abundance are highly variable between pink berry communities. Average read coverages for host (A) and phage (B) genomes were converted to reads per kilobase million (RPKM) using the total number of filtered and trimmed reads per sample.

### Pink berry phages are targeted by bacterial CRISPR systems

Bacterial CRISPR arrays serve as a record of past phage infection and can be used to infer hosts for phage genomes (Childs *et al*., 2014; England *et al*., 2018). Two independent CRISPR spacer prediction tools identified a total of 48 unique repeat sequences from four reference genomes for known pink berry-associated bacteria: *Desulfofustis* sp. PB-SRB1 (GenBank: JAEQMT010000010.1), *Flavobacteriales* bacterium (GenBank: DNTB01000031.1), *Rhodobacteraceae* sp. A2 (GenBank: JAERIM010000001.1), and *Thiohalocapsa* sp. PB-PSB1 (GenBank: CP050890.1) (Wilbanks *et al*., 2014). Parsing the remaining available pink berry genome, *Oceanicaulis alexandrii* sp. A1 (GenBank: JAERIO010000015.1), did not yield any CRISPRs. Because CRISPR repeat sequences are conserved within bacterial species (Lange *et al*., 2013; Mojica *et al*., 2000), they can be used to identify adjacent spacer sequences in unassembled metagenomic short reads (England *et al*., 2018; Skennerton *et al*., 2013). Using the CRISPR repeat sequences from reference genomes, NARBL (England *et al*., 2018) identified 2,802 unique CRISPR spacer sequences from the set of merged metagenomic reads from LS06-2018-s01, LS06-2018-s02, and LS06-2018-s03. Of these, 798 spacers were adjacent to repeats associated with *Desulfofustis* sp. PB-SRB1, 71 were adjacent to *Flavobacteriales* repeats, 349 were adjacent to *Rhodobacteraceae* sp. A2 repeats, and 1,584 unique spacers were adjacent to repeats associated with *Thiohalocapsa* sp. PB-PSB1.

To look for evidence of previous phage-bacteria interactions, metagenomic CRISPR spacer sequences were aligned to the eight complete phage genome assemblies (Fig. 1 & 4A). Of the 2,802 unique spacer sequences extracted from the merged set of metagenomic reads, 163 unique spacers aligned to seven phage contigs of interest with at least 80% identity over the entire spacer length (Fig. 1 & 4A, Suppl. Data 3). Spacers from three out of the four potential host taxa aligned to phage genomes, while no spacers from *Flavobacteriales* aligned to any phage genome of interest. *Thiohalocapsa* sp. PB-PSB1 was predicted to be the host of Thiohalocapsa phages MD03 and MD04, *Rhodobacteraceae* sp. A2 was predicted to be the host of Rhodobacteraceae phages MD05, MD06, and MD07, and *Desulfofustis* sp. PB-SRB1 was predicted to be the host of Desulfofustis phages MD01 and MD02 (Fig. 4A, Suppl. Data 3). For the two most prevalent and abundant phages identified, Desulfofustis phage MD02 (Suppl. Fig. 3A) and Thiohalocapsa phage MD04 (Suppl. Fig. 3B), phage genome and targeting CRISPR spacer coverages were positively correlated. Moreover, although most host spacers matched to a single virus, two spacers from the *Rhodobacteraceae* host aligned to Thiohalocapsa phage MD04 and Desulfofustis phage MD02 (Fig. 4A). We do not predict these phages to have infected the *Rhodobacteraceae* bacterium, since there was only one alignment each to Thiohalocapsa phage MD04 and Desulfofustis phage MD02 compared to 126 and three alignments from the two phage genomes to *Thiohalocapsa* and *Desulfofustis* spacers, respectively (Suppl. Data 3). Additionally, the alignments from the two *Rhodobacteraceae* spacers to Desulfofustis phage MD02 were weaker than the alignments from the *Desulfofustis* spacers (Fig. 4A, Suppl. Data 3). Alignments to multiple host taxa may be due to CRISPR systems targeting motifs that are present in several phage lineages.

**Figure 4.**
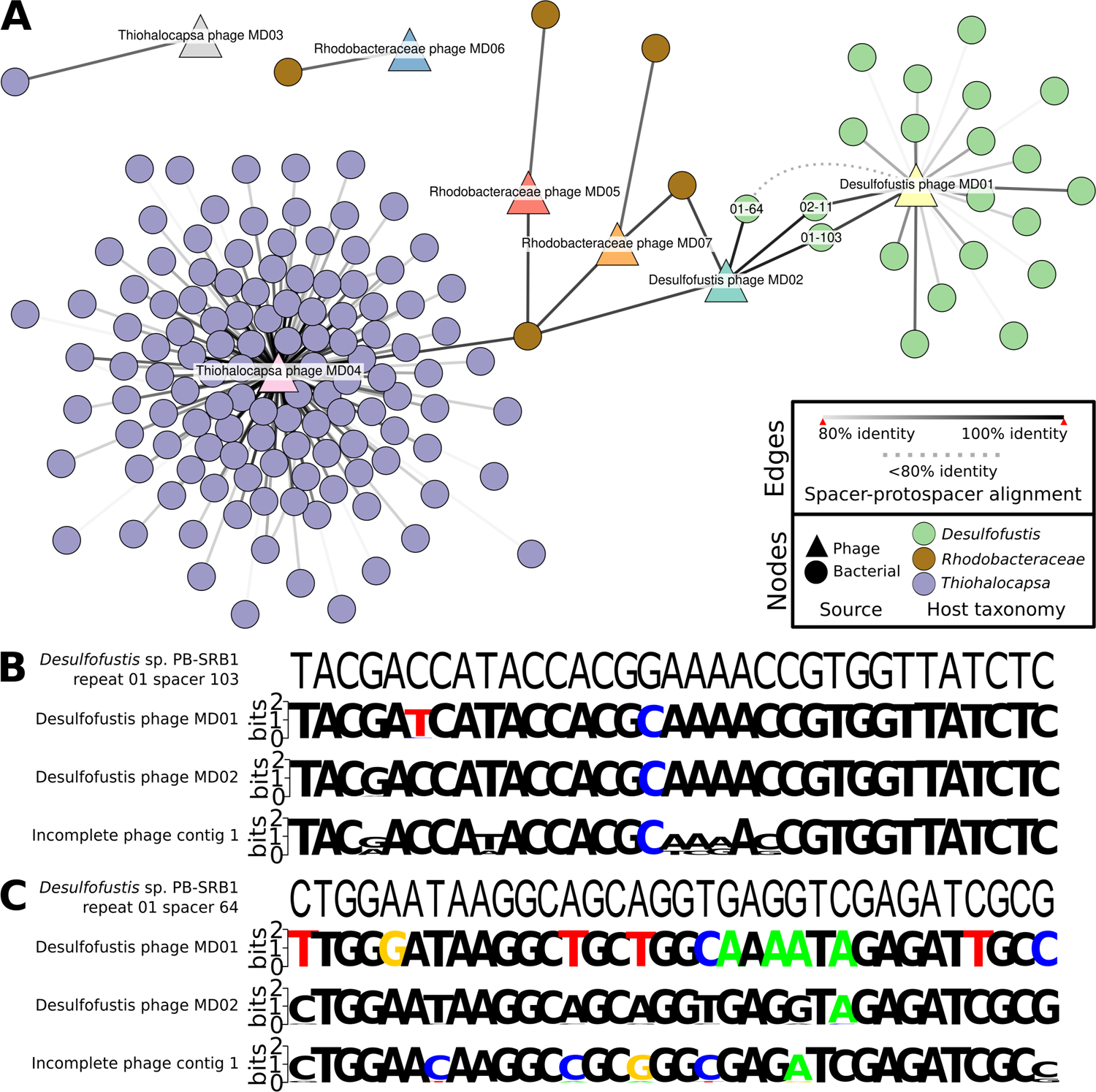
CRISPR spacer-phage genome alignments reveal hosts and a conserved protospacer. (A) Spacerblast (Collins & Whitaker, 2022) alignment results with at least 80% nucleotide identity over the entire spacer length were visualized in Cytoscape v3.9.0 (Su *et al*., 2014). Circular nodes represent unique spacers from bacterial contigs and are colored by taxonomy. Triangular nodes are phage contigs. Solid edges represent percent nucleotide identity over the entire spacer length. The dashed edge shows the connection between Desulfofustis phage MD01 and *Desulfofustis* sp. PB-SRB1 repeat 01 spacer 64, which is below 80% identity and included in part (C). The nucleotide sequences of phage protospacers within conserved capsid genes that were identified on phage contigs were aligned to (B) *Desulfofustis* sp. PB-SRB1 repeat 01 spacer 103 and (C) *Desulfofustis* sp. PB-SRB1 repeat 01 spacer 64. Spacer 02-11 is the reverse complement of 01-103 and is not shown. The resulting metagenome-wide variation in protospacer sequences from mapping reads to protospacers are shown as sequence logos (Crooks *et al*., 2004; Schneider & Stephens, 1990).

The vast majority of CRISPR spacer-protospacer matches occurred only once in the dataset. However, four spacers from *Thiohalocapsa* aligned imperfectly to two distinct protospacers on the genome of Thiohalocapsa phage MD04 (*Thiohalocapsa* sp. PB-PSB1 spacers 09-2, 21-2, 21-11, and 21-164 in Suppl. Data 3). An additional two spacers from *Desulfofustis* sp. PB-SRB1 are reverse complements of each other and target the same protospacer sequence on both Desulfofustis phage MD01 and MD02 (*Desulfofustis* sp. PB-SRB1 spacers 01-103 and 02-11) (Figs. 1, 4A, 4B, Suppl. Data 3). The shared CRISPR targeting of Desulfofustis phage MD01 and MD02 occurred in a conserved gene predicted to encode a phage capsid protein (Fig. 1, Suppl. Data 1) (ORFs DPMD01_45 and DPMD02_11, respectively), and corresponding to genomic regions with high read coverage relative to the remainder of the phage genomes (Fig. 1, Suppl. Fig. 2). A third spacer (01-64) targets a distinct protospacer within this same capsid gene on Desulfofustis phage MD02. A third capsid gene from an incomplete viral contig was found to be homologous to these two variants from Desulfofustis phage MD01 and MD02 and is 92% identical and of similar length. The capsid gene from this incomplete phage contig aligns with the same three spacers targeting Desulfofustis phage MD01 and MD02 (Fig. 4B, Suppl. Data 4). Nucleotide variation at these protospacers inferred by read mapping shows that other variants of these protospacers likely exist in related pink berry phages not assembled here (Fig. 4B). We obtained metagenome-wide allelic variants spanning the entire capsid gene of Desulfofustis phage MD01, Desulfofustis phage MD02, and Incomplete phage contig 1 and observed a near three-fold increase in the number of variants over CRISPR-targeted regions (Suppl. Fig. 4). Taken together, these results suggest that this conserved phage capsid gene is an active site of diversification.

### Horizontal gene transfer between pink berry phages and their hosts

Desulfofustis phage MD02 and Thiohalocapsa phage MD04 were found to contain discrete regions of high read coverage compared to the rest of the genome (Fig. 1, Suppl. Fig. 2). We hypothesized that these regions may be the result of read mapping from homologous regions of bacterial chromosomes or other phage genomes.

The high coverage region on Desulfofustis phage MD02 (genome coordinates 170-1760) did not align to any region of the *Desulfofustis* sp. PB-SRB1 host reference genome (GenBank: JAEQMT000000000.1) or to any other bacterial genomes in RefSeq. This region also did not successfully align to any other contigs in the co-assembly, indicating that the coverage at this region is not the result of conservation of this sequence among other members of the metagenome. Because of the circularity of these genomes, it is possible that the high read coverage at these regions is attributed to terminal redundancy from circular permutation (Garneau *et al*., 2017; Grossi *et al*., 1983).

The high coverage region of Thiohalocapsa phage MD04 (genome coordinates 21,035-22,077) encodes a glucosaminidase domain-containing protein (ORF TPMD04_36) predicted to function as the phage lysin. TPMD04_36 is homologous to two predicted ORFs (NCBI N838_07070 and N838_07065) in the *Thiohalocapsa* sp. PB-PSB1 genome (GenBank CP050890.1), which are adjacent to a predicted transposase (Fig. 5). This observation prompted an investigation into a possible transposon-mediated HGT event between Thiohalocapsa phage MD04 and its host, *Thiohalocapsa* sp. PB-PSB1.

**Figure 5.**
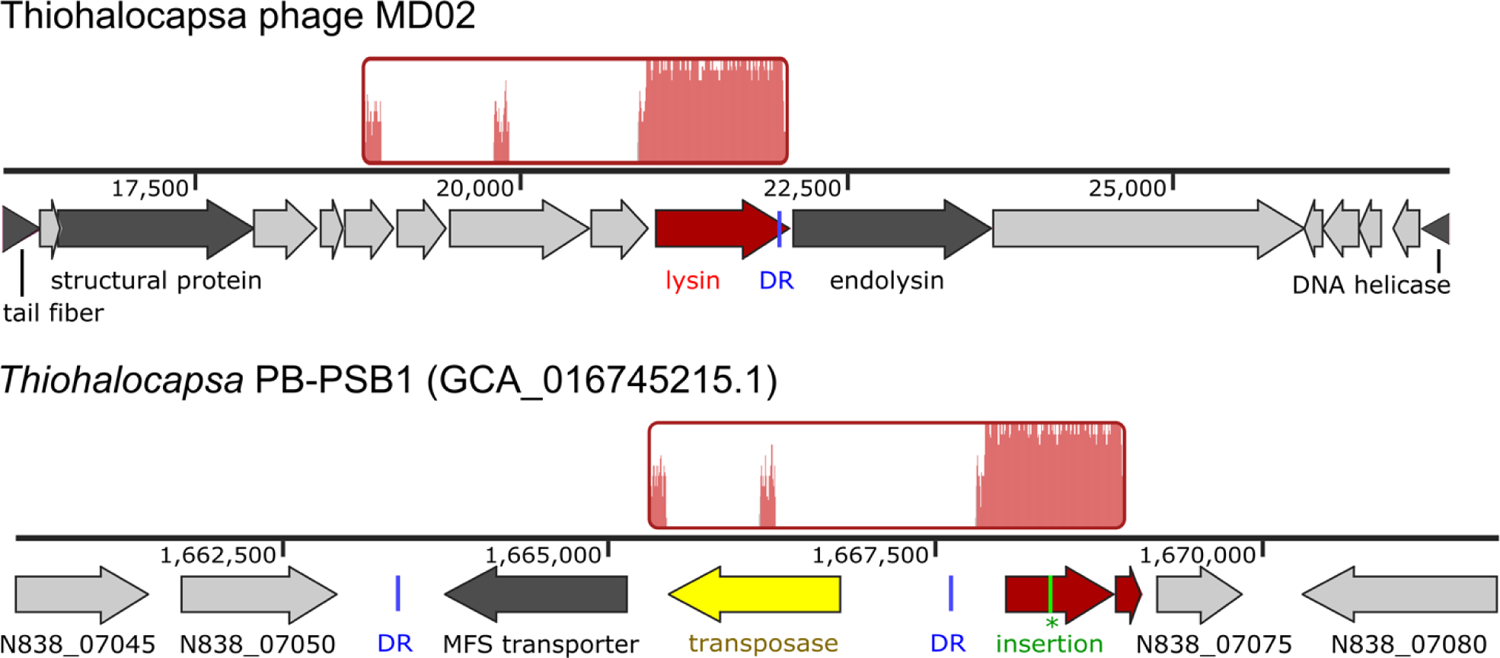
Region of homology between Thiohalocapsa phage MD04 and its host. Locally colinear blocks aligned with Mauve (Darling *et al*., 2004) are shown in red, with traces inside representing nucleotide similarity. Genome tracks show genome coordinates and ORFs. Conserved 17-bp direct repeat sequences (DR) are shown in blue. A single-nucleotide insertion (green) in the *Thiohalocapsa* ORF N838_07070 results in a premature stop codon and a pseudogene annotation.

Alignment of the Thiohalocapsa phage MD04 and *Thiohalocapsa* PB-PB1 genomes revealed that the phage lysin gene and the host pseudogene are in frame with each other, except the host ORF N838_07070 contains a single-nucleotide insertion at position 1,668,396 that results in a premature stop codon (Fig. 5). After closer observation of the region surrounding the pseudogene on the *Thiohalocapsa* sp. PB-PSB1 genome, we observed a transposon of the IS4 family with terminal inverted repeats and numerous direct repeats indicative of past transposase activity (Suppl. Fig. 5). The Thiohalocapsa phage MD04 genome contains an imperfect copy of a 17-bp direct repeat, differing by only one nucleotide, at the N-terminal of the TPMD04_36 ORF. Though MD04 was frequently targeted by host CRISPRs (including at protospacers directly adjacent to the lysin gene), no spacer-protospacer alignments were observed within the lysin gene, likely the result of selection against CRISPR self-targeting. Together, this finding suggests a past transposon-mediated HGT event may have resulted in the transfer of the phage lysin gene from Thiohalocapsa phage MD04, or a related ancestral phage, to its host.

## DISCUSSION

Marine phages are the most numerous biological components of the global ocean, outnumbering their bacterial hosts by tenfold (Breitbart *et al*., 2018), and play vital ecosystem roles as predators that turn over organic matter through bacterial lysis (Heldal & Bratbak, 1991; Maranger & Bird, 1995; Proctor *et al*., 1988; Steward *et al*., 1996) and as agents of HGT impacting bacterial community structure and function (Anantharaman *et al*., 2014; Breitbart *et al*., 2018; Kieft *et al*., 2021; Tuttle & Buchan, 2020). Pink berries are marine microbial aggregates with a microscale sulfur cycle, and have been used as a model system to study cryptic biogeochemical cycling (Wilbanks *et al*., 2014). Though phages have been identified within the pink berry metagenomes (Wilbanks *et al*., 2022), the full diversity of phages and their impacts on pink berry communities remain largely unexplored. Here, we investigated these simple, naturally-occurring microbial communities as a model for phage-host co-evolution.

We co-assembled three pink berry samples, recovering eight complete phage genomes spanning a total of 350 Kb and infecting three different bacterial species within the consortia, *Desulfofustis* sp. PB-SBR1, Rhodobacteraceae sp. A2, and *Thiohalocapsa* sp. PSB1. We found that pink berry-associated phages are highly diverse and largely novel, as seven of the eight complete phage genomes analyzed fail to cluster with any known phage sequences. One pink berry phage, Desulfofustis phage MD02, is only the second member of the genus *Knuthellervirus*. The other member of the *Knuthellervirus*, Pseudomonas phage PMBT14, infects *Pseudomonas fluorescens*, another marine organism, suggesting this phage genus infects diverse hosts.

Although the composition of the pink berry bacterial community is similar across individual aggregates, we found that phage presence and abundance is highly heterogeneous across samples. Pink berries are free-living microbial aggregates that exist at the sediment-water interface of intertidal ponds, with no obvious physical barrier to phage entry into the system. This raises ecological questions about the mechanisms underlying phage ingress into a pink berry aggregate and their persistence within, loss, or exclusion from the community.

CRISPRs are a common phage-resistance system employed by bacteria and archaea (Barrangou *et al*., 2007; Horvath & Barrangou, 2010; Jansen *et al*., 2002). We identified an astounding 2,731 unique CRISPR spacer sequences from pink berry-associated *Desulfofustis*, *Rhodobacteraceae*, and *Thiohalocapsa* hosts, 163 of which (~6%) target a complete phage genome we assembled. This discrepancy suggests that the true diversity of phages pink berry-associated bacteria encounter is far greater than what we report here. Seven of the eight phages investigated were targeted by described pink berry bacterial CRISPR systems, and the eighth, Pink berry phage MD08, may infect other pink berry-associated taxa that do not encode CRISPR defenses or for which we do not yet have a high-quality reference genome. Finally, we were able to observe diversification of a CRISPR-targeted conserved capsid gene, which is inconsistent with diversification across the rest of the phage genomes. Although we cannot establish a causative relationship between CRISPR targeting and phage variation, these observations are consistent with a model of CRISPR-driven evolution causing positive selection in a phage structural protein. We further observed a positive correlation between CRISPR spacer abundance and target phage abundance for two of the most prevalent and abundant phages identified. This suggests that these CRISPR spacers are positively selected for within individual pink berry consortia, and is consistent with observations from diverse microbial systems (Somerville *et al*., 2022; Meaden *et al*., 2021).

Phages and other mobile genetic elements are powerful mediators of HGT (Breitbart *et al*., 2018; Hall *et al*., 2017; Schneider, 2021). HGT between bacterial species within the pink berry consortia has been previously reported (Wilbanks *et al*., 2022), yet the role of phages in HGT and bacterial genome evolution in this system remains to be explored. We identified a predicted phage lysin gene that was horizontally transferred from Thiohalocapsa phage MD04 to its *Thiohalocapsa* host likely via a transposon intermediary. It is unclear if the lysin gene encoded on the *Thiohalocapsa* sp. PB-PSB1 genome is functional; a nonsense mutation in this ORF suggests it is a pseudogene, and was perhaps selected for to avoid deleterious effects of expressing this potentially lethal protein. Future work should aim to experimentally determine how these phages impact the evolution of both individual bacterial hosts and entire pink berry aggregates through HGT.

Taken together, our results demonstrate that pink berry communities contain diverse and variable phage consortia, which are highly targeted by host-encoded CRISPR systems. We leveraged metagenomic sequence data to better understand phage-host co-evolution occurring through CRISPR evasion and HGT. Pink berries offer a simple yet relatively unexplored, naturally-occurring model of phage invasion into and exclusion from microbial communities. The potential roles of phages in pink berry syntrophy and community-wide metabolic exchanges remain to be explored, but it is now clear that phages are notable members of these microbial consortia.

## METHODS

### Sampling

Pink berries, their surrounding sediment, and seawater were collected from pond LS06 (41.57587, −70.63781) in the Little Sippewissett Salt Marsh in Falmouth, MA, on July 17, 2018, using sterile 50-mL conical tubes. Samples LS06-2018-s01 and LS06-2018-s02 each contained one pink berry aggregate, and sample LS06-2018-s03 contained three pink berry aggregates. These samples were transported to the lab and immediately processed for DNA extraction.

### DNA isolation and sequencing

Pink berry samples were each mechanically homogenized in 1 mL of TE buffer and centrifuged at 1,000 x*g* for 1 min to pellet particulate matter. The supernatant was removed and subjected to a Wizard Genomic DNA Purification Kit (Promega Catalog No. A1120) according to the manufacturer’s instructions. Purified DNA was fragmented, and sequencing adapters and barcodes were ligated with the Nextera DNA Flex Library Prep Kit (Illumina Catalog No. 20018705) using Nextera DNA CD indexes (Illumina Catalog No. 20018708). DNA yield was measured with a Qubit High Sensitivity dsDNA assay kit (ThermoFisher Catalog No. Q32851), and DNA from LS06-2018-s01, LS06-2018-s02, and LS06-2018-s03 were pooled at a ratio of 1:1:10. After pooling, DNA was purified with Ampure XP beads (Beckman Coulter Catalog No. A63881) according to the manufacturer’s instructions. DNA sequencing was performed on an Illumina HiSeq 2500 using a 2×250nt protocol at the University of Illinois at Urbana-Champaign Roy J. Carver Biotechnology Center.

### Metagenome co-assembly & read mapping

For each sample, reads were quality checked with FASTQC v0.11.9 (Andrews, 2010), trimmed using Trimmomatic v0.39 (Bolger *et al*., 2014), and adapter sequences were removed. Trimmomatic filtered reads using a sliding window of 4 base pairs, a minimum average base quality score of 15, a minimum quality score for retention on the leading and trailing ends of 2, and a minimum read length of 100 bases. The resulting trimmed and filtered reads were merged into a single set of reads and co-assembled using Metaviral SPAdes v3.15.2 (Antipov *et al*., 2020) with default parameters to maximize the recovery of complete, circular phage genomes. To estimate phage and bacterial abundances, Bowtie2 v2.4.5 (Langmead & Salzberg, 2012) was used to map reads from the set of merged reads or from each pink berry sample to assembled phage contigs and to representative host genomes from NCBI BioProject PRJNA684324: *Desulfofustis* sp. PB-SRB1 (GenBank: JAEQMT010000010.1), *Rhodobacteraceae* sp. A2 (GenBank: JAERIM010000001.1), and *Thiohalocapsa* sp. PB-PSB1 (GenBank: CP050890.1) (Wilbanks *et al*., 2022). Read mapping statistics were obtained from Bowtie2 alignments using samtools v1.15.1 (Danecek *et al*., 2021).

### Phage sequence identification, binning, and annotation

ViralVerify v1.1 (Antipov *et al*., 2020) was used with default settings to categorize the metagenome contigs as putatively bacterial or viral. ViralComplete v1.1 (Antipov *et al*., 2020) was used with default settings to identify viral contigs that represent complete phage genomes. To verify these predictions, VIBRANT v1.2.1 (Kieft *et al*., 2020) was used with default settings on the same metagenome contigs. All resulting putative viral contigs that were estimated to be complete viral genomes by both prediction tools were targeted for downstream analyses and annotation. vConTACT v2.0 (Bin Jang *et al*., 2019) was used with the RefSeq v211 Viral database (Brister *et al*., 2015) to cluster the viral contigs of interest with existing phage genomes and to approximate phage taxonomy.

The viral contigs of interest were passed through Pharokka v1.0.1 (github.com/gbouras13/pharokka), using PHANOTATE v1.5.0 (McNair *et al*., 2019) to predict genes and PHROGs v3 (Terzian *et al*., 2021) to provide initial protein annotations. Protein functions were also predicted using Phyre2 (Kelley *et al*., 2015), BLASTp v2.11.0 (Altschul *et al*., 1990) with the NCBI RefSeq v211 virus amino acid database and the non-redundant amino acid database (Brister *et al*., 2015; O’Leary *et al*., 2016), and HMMER v3.2.1 (Eddy, 2011) with Pfam-a v35.0 (Mistry *et al*., 2021) and TIGRFAMs v15.0 (Li *et al*., 2021). The resulting predictions from each method were manually reviewed for each protein, and a consensus annotation was inferred (Suppl. Data 1). Clinker v0.0.23 (Gilchrist & Chooi, 2021) was used to identify conserved genes among phage genomes, which were then aligned with tBLASTx v2.11.0 (Camacho *et al*., 2009) against all the contigs in the metagenome co-assembly. Metagenomic reads from pink berries sampled in 2011 (SRA: SRR13297012) were mapped to the viral contigs of interest with Bowtie2 using the same methods described above.

### CRISPR spacer-protospacer analysis and host prediction

CRISPRclassify v1.1.0 (Nethery *et al*., 2021) and MinCEd v0.4.2 (Bland *et al*., 2007) were used to identify CRISPR repeat sequences from representative genomes of pink berry taxa from NCBI BioProject PRJNA684324: *Desulfofustis* sp. PB-SRB1 (GenBank: JAEQMT010000010.1), *Oceanicaulis alexandrii* (GenBank: JAERIO000000000.1), *Rhodobacteraceae* sp. A2 (GenBank: JAERIM010000001.1), *Thiohalocapsa* sp. PB-PSB1 (GenBank: CP050890.1) (Sayers *et al*., 2020; Wilbanks *et al*., 2022), and *Flavobacteriales* bacterium (GenBank: DNTB01000031.1). The identified repeats from each tool were combined and dereplicated to obtain a list of repeats found in the genomes of pink berry taxa. Since CRISPR arrays are often misassembled with short-read data, putative spacer sequences from the trimmed and filtered reads of each LS06-2018 metagenome were identified with NARBL (England *et al*., 2018) using the dereplicated set of repeats identified from the reference genomes, an approximate repeat size of 36, and a minimum coverage of supporting neighbor spacers of 2. Since repeat sequences are highly conserved between bacterial species (Kunin *et al*., 2007), any spacer identified by NARBL was inferred to belong to the same species as the reference genome from which its associated repeat came. The resulting spacers were aligned to the viral contigs of interest using Spacerblast v0.7.7 (Collins & Whitaker, 2022). Viral contigs that aligned to spacers with at least 80% identity over the full length of the spacer were considered to have a host match with the bacterium whose genome contained the spacer. The merged set of filtered and trimmed reads were aligned to spacers identified by NARBL with Bowtie2 and read coverage statistics were obtained with samtools as described above.

Metagenome reads were mapped to the regions containing conserved phage genes identified with Clinker or tBLASTx, above, with spacer-protospacer alignments with Bowtie2. Mapped reads were converted to multiple sequence alignments using SAM4WebLogo in JVarkit v2021.10.13 (Lindenbaum, 2015) and sequence logos were visualized with WebLogo (Crooks *et al*., 2004). Allele variants for conserved phage genes were called by using Snippy v3.2 (github.com/tseemann/snippy) with default settings on metagenome reads. Variants that resulted before the “--mincov” and “--minfrac” filters were applied were used downstream to maximize the number of possible variants recovered. Variant statistics were obtained and visualized using vcfR v1.13.0 (Knaus & Grünwald, 2017) with default settings, except 100-bp size windows were used instead of 1000-bp.

### Horizontal gene transfer analysis

Uneven sequence coverage patterns on phage genomes are sometimes attributed to HGT between the phage and its host genome, especially if the region aligns to the host genome and/or contains genes that facilitate HGT (Kleiner *et al*., 2020). Viral genomes of interest and their coverages from the merged set of metagenome reads were visualized in IGV v2.11.4 (Thorvaldsdóttir *et al*., 2013). For any discrete regions of the viral genomes of interest that had much higher read coverage (>3x) than the surrounding region, those regions were aligned to their predicted host genomes, if available, using BLASTn v2.11.0 (Camacho *et al*., 2009). Predicted regions of HGT between phages and their hosts were aligned with Mauve v2015-02-13 (Darling *et al*., 2004). Conserved repeats flanking a putative transposon were initially identified by reciprocal BLASTn using “blastn-short”. The transposon region with its inverted repeats in the *Thiohalocapsa* sp. PB-PSB1 genome was identified with ISEScan v1.7.2.3 (Xie & Tang, 2017), and other repeats were identified and annotated manually in Geneious Prime v2022.1.1 (www.geneious.com/prime).

## Data availability

DNA sequencing reads from this study are deposited in the NCBI SRA under PRJNA907316. Assembled phage genomes are deposited in NCBI GenBank under the following accession numbers: OP947158.1 (Desulfofustis phage MD01), OP947159.1 (Desulfofustis phage MD02), OP947165.1 (Thiohalocapsa phage MD03), OP947166.1 (Thiohalocapsa phage MD04), OP947161.1 (Rhodobacteraceae phage MD05), OP947162.1 (Rhodobacteraceae phage MD06), OP947163.1 (Rhodobacteraceae phage MD07), OP47164.1 Pink berry phage MD08, OP947160.1 (Pink berry virus MD00).

## ACKNOWLEDGMENTS

This research was conducted as a part of the 2020 Microbial Diversity course at the Marine Biological Laboratory (MBL), which was funded by NSF Award ID 1822263, DOE Award ID DE-SC0016127, and the Simons Foundation. We gratefully acknowledge the insights and support provided by all participants and instructors, especially Whitney England, Laura Suttenfield, and George O’Toole. D.E.C. was supported by NIH T32 DK077653-29 and Crohn’s & Colitis Foundation Research Fellowship Award #935619. E.G.W. gratefully acknowledges support from the Whitman Fellowship at the MBL. This research was sponsored by the U.S. Army Research Office and accomplished under cooperative agreement W911NF-19-2-0026 for the Institute for Collaborative Biotechnologies.

Conceptualization, J.C.K., D.E.C., E.G.W., and R.J.W.; Data Curation, J.C.K., D.E.C. and R.J.W.; Analysis, J.C.K. and D.E.C.; Funding Acquisition, R.J.W.; Investigation, J.C.K. and D.E.C.; Methodology, J.C.K., D.E.C., E.G.W., and R.J.W.; Project Administration, J.C.K., D.E.C., E.G.W., and R.J.W.; Resources, R.J.W.; Software, J.C.K. and D.E.C.; Supervision, D.E.C., E.G.W., R.J.W.; Validation, J.C.K. and D.E.C.; Visualization, J.C.K. and D.E.C.; Writing—Original Draft, J.C.K. and D.E.C.; Writing—Review and Editing, J.C.K., D.E.C., E.G.W., and R.J.W.

