## Supplemental figures for "Horizontal gene transfer and CRISPR targeting drive phage-bacterial host interactions and co-evolution in pink berry marine microbial aggregates"

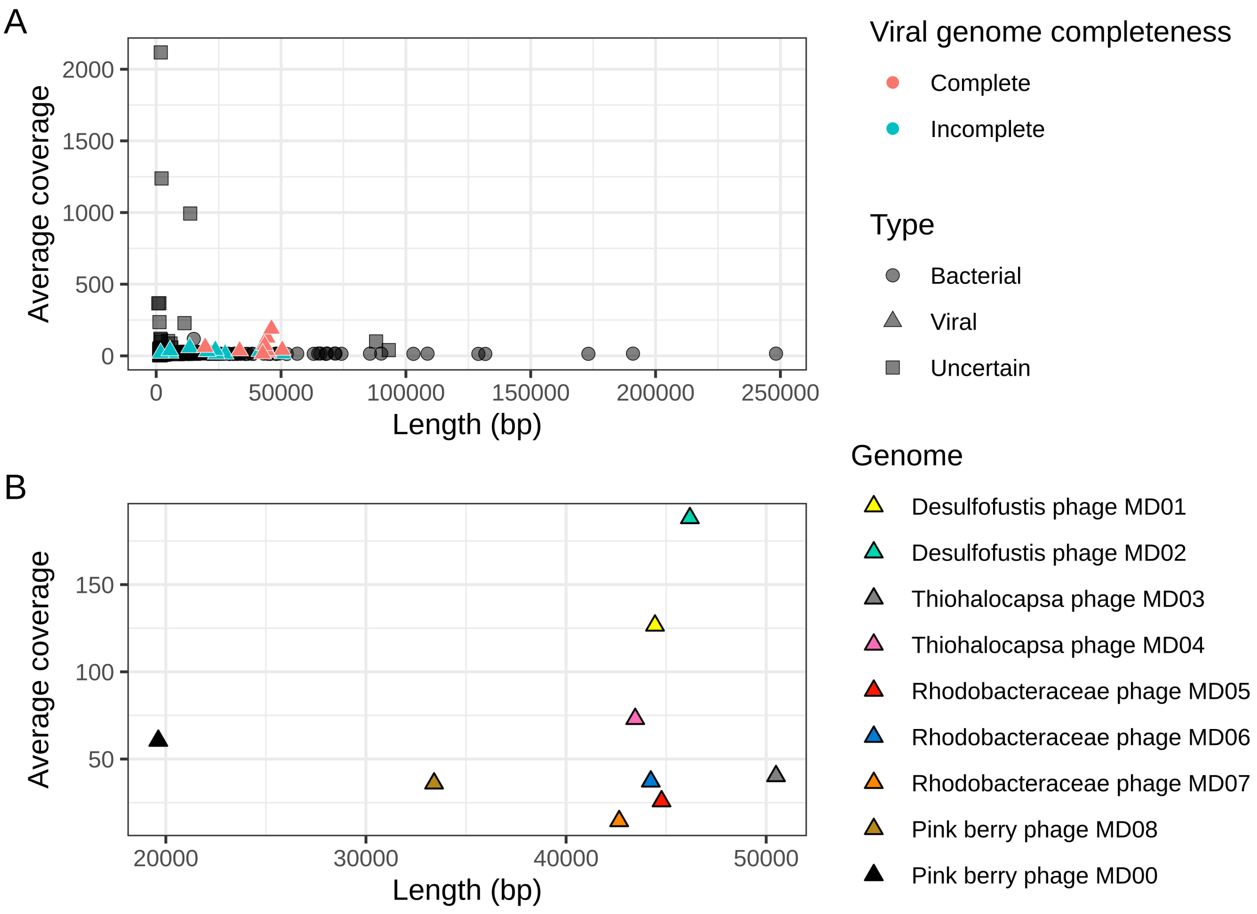


**Supplemental Figure 1. Coverage and length of co-assembled pink berry contigs.** Contig types, and viral genome completeness were obtained from ViralVerify (Antipov *et al.*, 2020) and ViralComplete (Antipov *et al.*, 2020), respectively. Coverage values were calculated using the merged set of filtered and trimmed reads used for metagenome co-assembly. (A) All contigs from Metaviral SPAdes coassembly, (B) the nine complete, viral verified contigs which were of interest for downstream analyses.

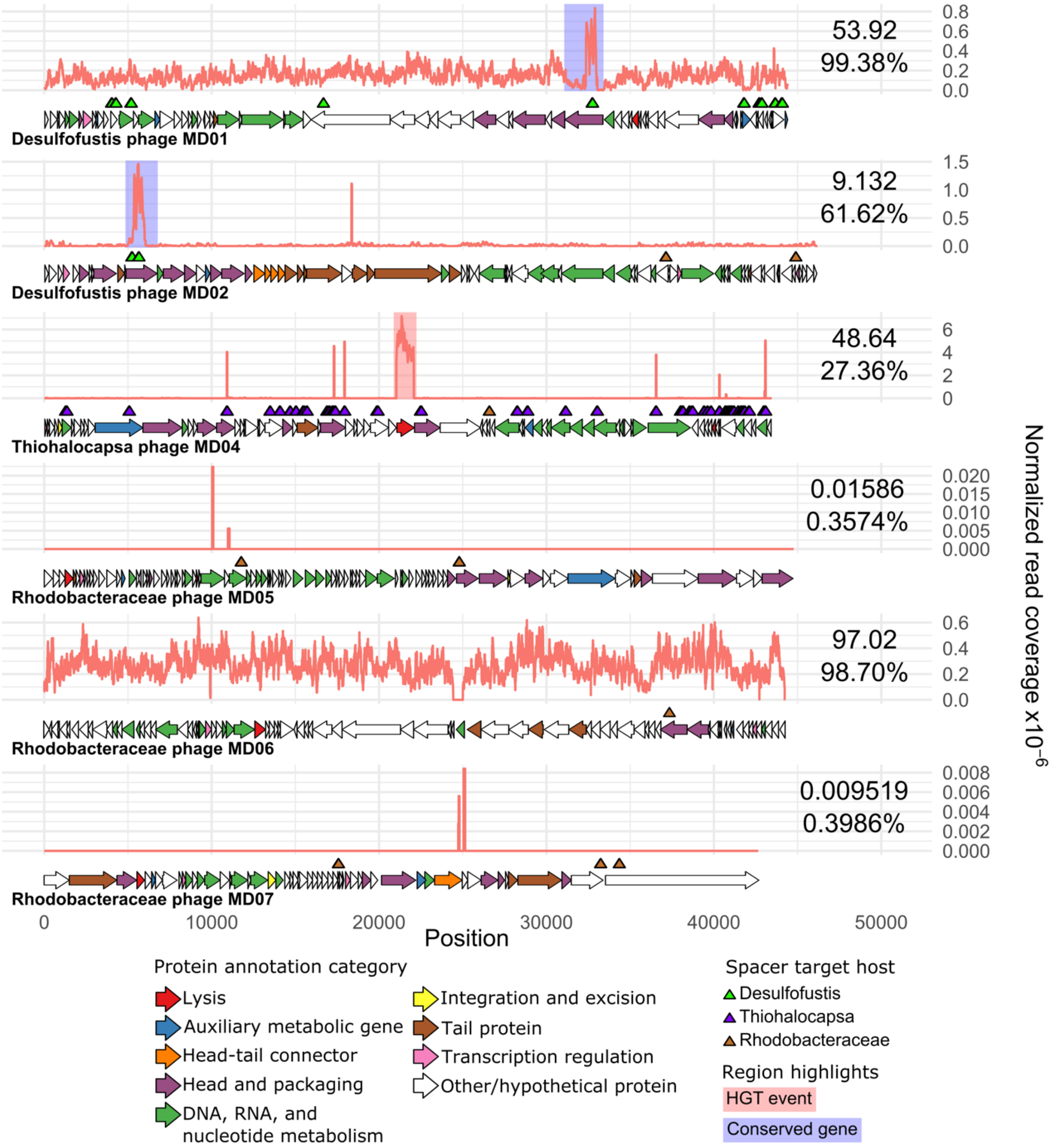


**Supplemental Figure 2.** Normalized read coverage by position (orange lines) from NCBI SRR13297012, representing a metagenome of a pool of 10 pink berries sampled in 2011. Reads from this sample did not map to Thiohalocapsa phage MD03 or Pink berry phage MD08, which are not shown. Values for the non-normalized average read coverage and the percent of the genome that is covered are shown on the right for each genome. Coverage values were normalized by dividing the depth at a given position by the total number of reads in sample SRR13297012. Horizontal arrows indicate ORFs predicted by PHANOTATE (McNair *et al.*, 2019), and their colors correspond to predicted functional categories. Triangles indicate genome positions of protospacers, colored by host taxonomy of the corresponding spacer. Regions highlighted with a blue background indicate a conserved gene between viral genomes inferred by Clinker (Gilchrist & Chooi, 2021). Region highlighted with a red background was found to be an HGT event between phage and host.

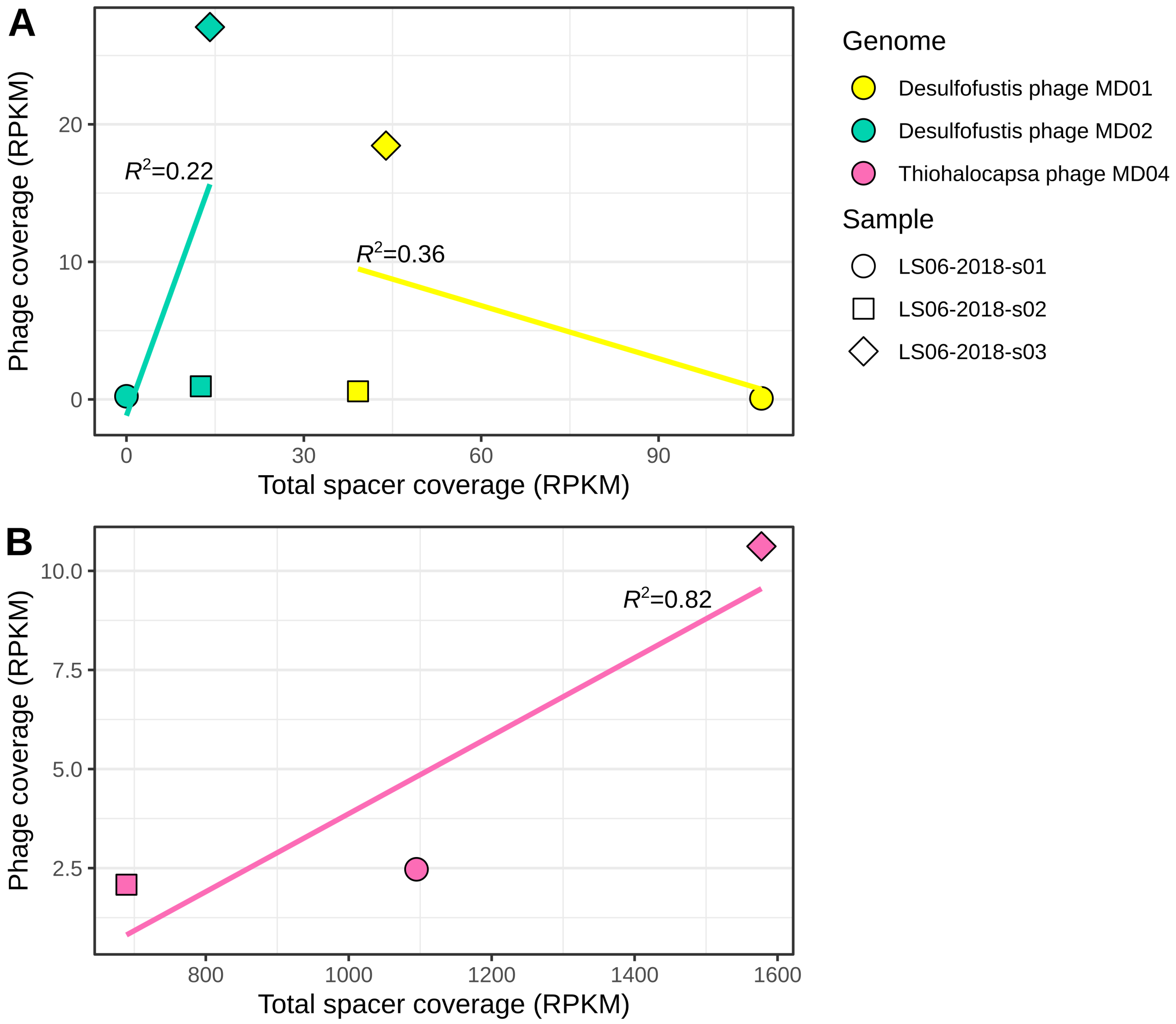


**Supplemental Figure 3.** Average read coverages for CRISPR-targeted phage genomes and spacer sequences were converted to reads per kilobase million (RPKM) using the total number of filtered and trimmed reads per sample. For a given phage genome, the RPKM for all spacers known to target it were added to give the total spacer coverage. Thiohalocapsa phage MD03, Rhodobacteraceae phage MD05, Rhodobacteraceae phage MD06, and Rhodobacteraceae phage MD07 each had spacer coverages below detection and were excluded. Linear regressions were fitted for each phage genome and their associated multiple R-squared values are provided. (A)

Phage coverage versus total spacer coverage for Desulfofustis phage MD01 and Desulfofustis phage MD02. (B) Phage coverage versus total spacer coverage for Thiohalocapsa phage MD04.

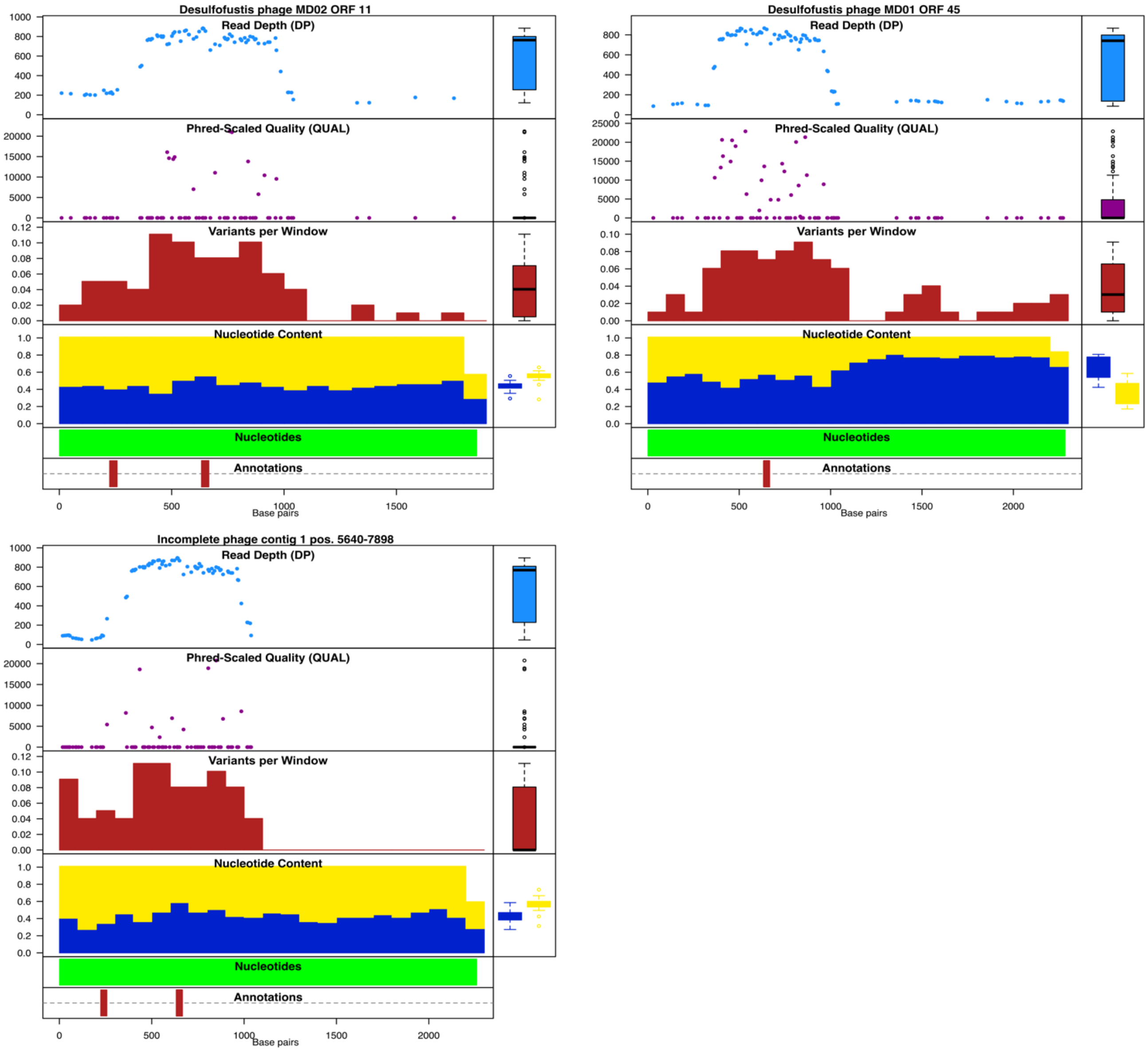


**Supplemental Figure 4**. Metagenome-wide variant calling spanning a conserved predicted capsid gene present in Desulfofustis phage MD01 (DPMD01_45), Desulfofustis phage MD02 (DPMD01_11), and Incomplete phage contig 1. Metrics were obtained using 100-bp windows. Read depth and mapping quality metrics are only provided for positions with an alternate variant. Regions targeted by a host CRISPR spacer are designated by red bars as Annotations.

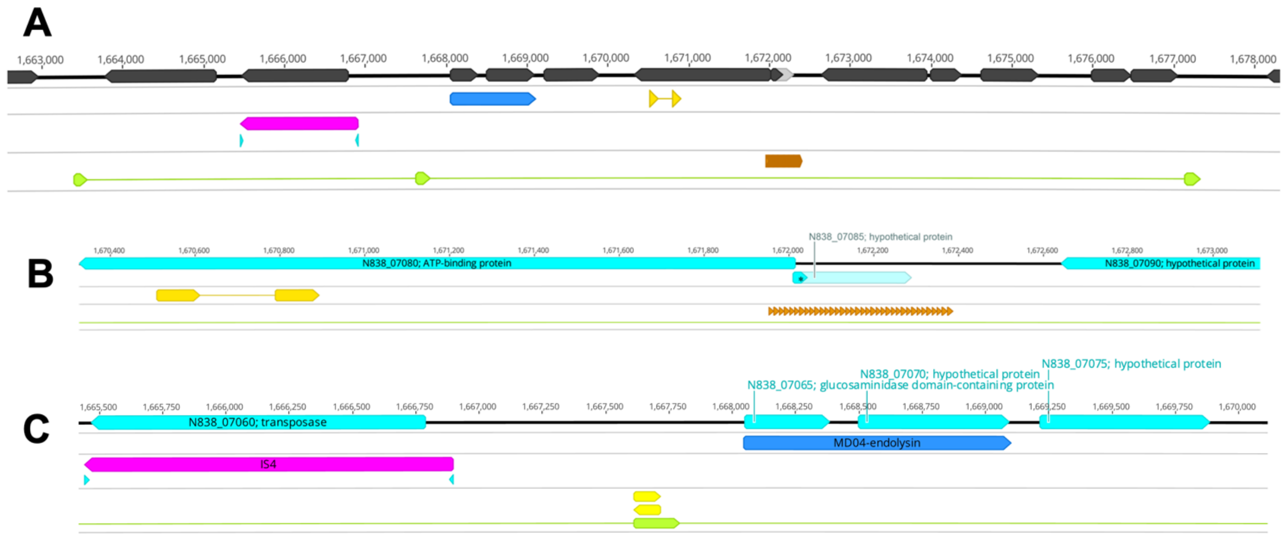


**Supplemental Figure 5**. The genomic region of the *Thiohalocapsa* sp. PB-PSB1 genome (GenBank: CP050890.1) surrounding the endolysin pseudogene region horizontally transferred from Thiohalocapsa phage MD04 (dark blue) contains a transposon of the IS4 family (pink) with terminal inverted repeats (light blue arrows) and numerous direct repeats indicative of past transposase activity. (A) In a 15-kilobase region surrounding the horizontally transferred region (dark blue arrow), we observed a 102 base pair direct repeat (orange arrows, HR01-HR02) that is also found in the Thiohalocapsa MD04 genome, an intergenic 183-bp direct repeat (green arrows, AR01-AR03), and a 13-bp imperfect tandem repeat region (brown arrows) that begin within an ATP-binding protein and extend through a neighboring pseudogene and intergenic region. (B) An enlarged view of the region containing this tandem repeat region (brown arrows) and the ATP-binding protein gene with the MD04-shared repeats (orange arrows) to clearly show the 35 tandem repeat units, which we have named QR01-QR35 (sequences provided in Suppl. Data 5). (C) An enlarged view of the region including the HGT region (dark blue) and the transposon (pink). The transposon, PSB1_IS134 was identified as a member of the IS4 family and is flanked by terminal inverted repeats. Here, we also show that the 183-bp direct repeats (green arrow), also contain an imperfect palindromic repeat sequence at their 5' ends (indicated by opposing yellow arrows). Sequences and genome coordinates for all these repeat regions are provided in Suppl. Data 5.
